# Navigating methodological decisions: Balancing rigor and data volume of the Canadian Living Planet Index

**DOI:** 10.1101/2025.09.15.676271

**Authors:** Jessica Currie, Sarah M Ravoth, Valentina Marconi, Louise McRae, Maria I Arce-Plata, Sandra Emry, Robin Freeman, Maximiliane Jousse, Gaëlle Mével, Shuaishuai Li, Cristian A Cruz-Rodríguez, David AGA Hunt, Philippa Oppenheimer, Lauren Gill, Janaina Serrano, Stefanie Deinet

## Abstract

The Living Planet Index — a biodiversity indicator that assesses the relative change of aggregate vertebrate abundance data — is an indicator used in global and national biodiversity monitoring frameworks. In Canada, the LPI has been modified (C-LPI) — adopting differing methodological choices relative to the global LPI. However, there is no clear consensus on the most appropriate analytical methods, particularly as they pertain to the treatment of zeros, confidence intervals and uncertainty, time series length and number of data points required, modelling of short time series, removal of outliers, weighting species, and the impact of baseline year selection. Our analysis transparently explores multiple methodological options and the consequent C-LPI output for each of these decision points. Our research does not evaluate the superiority of a single approach but rather provides transparency and accountability in C-LPI reporting. We hope that this will further strengthen the utility of the C-LPI and provide decision makers with the necessary information to appropriately interpret patterns, evaluate progress towards biodiversity targets, and inform conservation action.

## Introduction

Addressing biodiversity loss demands ambitious targets, rapid implementation and reliable indicators to accurately track progress towards national (e.g., Canada’s 2030 Nature Strategy) and global (e.g., Convention on Biological Diversity’s Kunming-Montreal Global Biodiversity Framework) biodiversity goals. The Living Planet Index (LPI) — conceived over 25 years ago — is a biodiversity indicator used to evaluate the state of wildlife by assessing the relative change of aggregate monitored vertebrate population abundance (Ledger *et al*. 2023; Loh *et al*. 2005). The indicator has been applied at global (WWF 2024), national (Marconi *et al*. 2021; Bayraktarov *et al*. 2020; WWF-Canada 2020), and regional scales (McRae *et al*. 2012) and has also garnered widespread uptake as a public communications tool (Ledger *et al*. 2023).

Originally conceived by the World Wildlife Fund, the LPI dataset and methods have been continually developed to advance accuracy, address potential biases, and appropriately report on average trends in relative vertebrate abundance (Ledger *et al*. 2023). In Canada, the LPI methodology has been adapted to produce a biodiversity indicator that supports domestic policy reporting (ECCC 2024). The indicator is coined the Canadian Species Index (CSI; ECCC 2023) by government agencies and is synonymous with the Canadian Living Planet Index (C-LPI; WWF-Canada 2020) — a title commonly used by WWF-Canada^1^

However, like other high-profile biodiversity indicators (e.g., IUCN Red List; Turnhout and Purvis 2021; Rodrigues *et al*. 2006), the LPI has garnered criticism in recent years, which has cascading impacts for domestic policy reporting. Criticisms are due, in part, to interpretation and communication issues (Ledger *et al*. 2023; Puurtinen *et al*. 2022), but also a result of questions arising from the underlying data and methods (Leung *et al*. 2022a; Puurtinen *et al*. 2022; Leung *et al*. 2020). The LPI relies on a geometric mean of relative abundance (Buckland *et al*. 2011; Collen *et al*. 2009), and while useful for its sensitivity to change (Santini *et al*. 2017), comprises statistical challenges where methodological decisions must be appropriately made, including how to address population counts of zero and potential outliers (Leung *et al*. 2020; Buckland *et al*. 2011). These decisions are not straightforward — each with its own merits and drawbacks. For instance, the global LPI applies 1% of the mean of the population time series to all values within the time series containing zero values (Collen *et al*. 2009), recognizing the need to incorporate potential population crashes and subsequent recoveries. However, when zeros are replaced with a non-zero number, the geometric mean can be sensitive to the value chosen (Buckland *et al*. 2011). Alternatively, the C-LPI/CSI treats zeros within the time series as missing values (Marconi *et al*. 2021), because on close inspection of these data, they rarely reflect species extirpations. Other methodological considerations and points of contention include assessments of uncertainty (Johnson *et al*. 2024), time series length and number of data points (Marconi *et al*. 2021), baseline values (Ledger *et al*. 2023), weighting of data (e.g., McRae *et al*. 2017), and modelling approaches (Ledger *et al*. 2023; Leung *et al*. 2020). Critically, however, there is no clear consensus on how to appropriately address these issues within the framework of the LPI.

We aim to demonstrate how different methodological choices influence the C-LPI outputs. Methodological choices include:

i. treatment of zeros (replacing zero values to permit mathematical calculation of the index);
ii. confidence intervals and uncertainty (understanding uncertainty associated with the index);
iii. time series length, number of data points required and completeness (evaluating criteria for data inclusion);
iv. modelling of short time series (assessing differing mathematical applications);
v. removal of outliers (evaluating the influence of extreme increases and declines) ;
vi. weighting species (assessing options for addressing species representation biases); and
vii. baseline year (understanding the influence of the baseline year on interpretation of trends).

These topics were selected because they have recently been the subject of criticism and are believed to influence the LPI. Specifically, we used the C-LPI/CS^2^ data as the underlying information offers some of the best representation of biodiversity, arising from the number of species (ECCC 2023; Marconi *et al*. 2021), as well as taxonomic breadth and distribution of biotic variables (Currie *et al*. 2022) that it includes. Our exploratory analysis emphasizes end points as they represent the current metric used in public reporting, while qualitatively noting trajectory differences or consistency, where visually apparent. Taken together, we qualitatively evaluate the sensitivity of the C-LPI to methodological decisions criticized within geometric relative abundance indicators more broadly. Because there is no agreed quantitative bench for the ‘true’ trajectory of wildlife populations, our research instead, aims to emphasise the importance of transparency and accountability and aid in the continued development of the LPI.

## Methods

### Data

Population time series data for Canada were obtained from the most recent Living Planet Index Database (LPD 2023) and were supplemented by additional data collection^3^ The population time series data were gathered from a variety of sources, predominantly government databases and peer-reviewed publications, following the criteria outlined by Collen *et al*. (2009). In alignment with domestic policy reporting (ECCC 2023), data were restricted to native vertebrate species that regularly occur in Canada, according to the 2020 Wild Species Report (CESCC 2022). Species classified as “Not Applicable”, “Presumed Extirpated” or “Probably Extirpated” were excluded from the dataset. In cases where there was spatial and temporal overlap of population time series for a given species, we retained only one of the overlapping populations to reduce geographic sampling bias (i.e., replicates were removed). Priority for inclusion was given to higher quality data, defined by greater time series length and number of data points, methods and credibility of the data source (e.g., priority given to long-term monitoring programs). These criteria produced a dataset comprising 5,793 monitored populations of 950 vertebrate species native to Canada. Despite its impressive representation (Currie *et al*. 2022, Marconi *et al*. 2021), the dataset comprises population time series containing varying lengths, data points and zero values. Moreover, there are different numbers of population time series and species contributing to each year of the index, and each index output investigated below.

### Methodological decisions

The LPI is calculated as the geometric mean of the relative changes in population abundance within a species (equivalent to averaging the logarithm of rate of change of abundance for each species), after which species-specific trends are aggregated using a geometric mean to generate a single annual index (see Loth *et al*. 2005 for details on the mathematical procedure). We showcase the impact of seven methodological decisions associated with the measurement of the C-LPI: (i) treatment of zero values, (ii) confidence intervals and uncertainty, (iii) time series length, number of data points and completeness, (iv) modelling of short time series, (v) removal of outliers, (vi) weighting of species, and (vii) selection of the baseline year. Methodological changes were compared to current approaches employed by the C-LPI (Table 1). Native vertebrate trends were calculated predominantly using selectable options under the publicly available *rlpi* R package (Freeman *et al*. 2017), with indices spanning from 1970 to 2022. The data and code for investigating indices are available through Github (https://github.com/ciee-ldp/CanadianLPI_shiny.git), though confidential records have been removed from the online repository.

**Table 1.** Current methodological approaches adopted by the global Living Planet Index (LPI) and the Canadian Living Planet Index (C-LPI).

| Methodological decision | Global LPI approach | C-LPI approach |
| --- | --- | --- |
| i) Treatment of zero values | Adding 1% of the mean of the population time series to all values within the time series containing zeros | Treated as missing values (i.e. replaced with NA) |
| ii) Confidence intervals and uncertainty | Bootstrapping by randomly sampling species for each year | Bootstrapping species trends for all years (10,000 bootstrap resamples) |
| iii) Time series length and number of data points<br>iv) Modelling of short time series | Requirement of 2 or more data points, no restriction on length<br>Log-linear interpolation on time series with <6 data points and a generalized additive model (GAM) on time series with ≥6 data points | Requirement of 3 or more data points, no restriction on length<br>Linear regression models on time series with <6 data points and a generalized additive model (GAM) on time series with ≥6 data points |
| v) Removal of outliers | No outliers routinely removed | No outliers routinely removed |
| vi) Weighting species | Species weighted by diversity | Species weighted equally |
| vii) Baseline year | 1970 | 1970 |

### (i) Treatment of zero values

Mathematically, population counts of zero pose a problem within the calculation of the LPI, as all values are logged in the process of calculating the geometric mean. Consequently, all population counts of zero must be replaced. The current C-LPI approach is to treat zeros as missing values (referenced as *NA (C-LPI)*) — in line with recommendations on the calculation of the index (Toszogyova *et al*. 2024). This decision was based on close investigation of the time series in the dataset, which were deemed to be more often missing observations rather than representations of extirpation (Marconi *et al*. 2021). Here, we explore six alternatives:

a. *+1% mean*: Adding 1% of the mean of the population time series to all values within the time series containing zeros (this approach is currently adopted by the global LPI; Collen *et al*. 2009).
b. *+minimum*: Adding the minimum non-zero value of the time series to all values within the time series containing zeros.
c. *+1*: Adding 1 to all values within the time series containing zeros.
d. *+0*.*000001*: Adding a small value (0.000001) to all zeros.
e. *NA, NA, +1% mean:* Conditionally replacing zero values depending on their position in the time series. Leading zeros (i.e., those at the beginning of the population time series) and zeros located in the middle (i.e., those with >0 values in years before and after the zero value) are treated as missing values, while trailing zeros (i.e., those at the end of the population time series) are kept (by adding 1% of the mean to all values within time series containing zeros, as in (a) on the assumption that they might represent local extirpations. This allows for inclusion of small values that approximate zero, rather than treating these potential extirpations as uninformative (i.e. replacement with NA).
f. *+1% mean, NA, +1% mean:* Conditionally replacing zero values depending on their position in the time series. Leading zeros and trailing zeros are kept by adding 1% of the mean to all values within time series containing zeros, on the assumption that they might represent, respectively, recolonizations and local extirpations, while zeros located in the middle are treated as missing values.

Options *a-d* are available in the *rlpi* package, while options *e* and *f* are novel explorations.

### (ii) Confidence intervals and uncertainty

Ideally, uncertainty would be propagated up from each population, but information on uncertainty for individual population counts is not included in the database and often differs among sources. Consequently, confidence intervals here reflect the range of LPI values that can fit into the existing data, capturing the variability within the dataset rather than the true variability associated with population counts, which would be reflected within a typical confidence interval. The C-LPI presents 95% confidence intervals calculated from bootstrapping logged interannual change values (“lambdas” hereafter) with replacement within *species* and produces a value for each of the 10,000 bootstrapped resamples. This approach aims to account for autocorrelations in lambdas over time for a given species. Here, we explore two alternatives using 10,000 bootstrap samples:

a. Adopting the global LPI default setting in the *rlpi* package (Freeman *et al*. 2017) to calculate confidence intervals around LPI trends by bootstrapping lambdas within a *year* and taking the 95% central values to construct the confidence interval values for that year. This approach was used by Collen *et al*. (2009), where bootstrapping was conducted by randomly sampling species for each year, independently, which did not account for autocorrelations in lambdas over time for a given species.
b. Bootstrapping lambdas within a *population* and re-averaging at the species level to produce 95% confidence intervals for the 10,000 bootstrapped resamples. This approach aims to account for autocorrelations in lambdas over time for a given population.

Option *a* is available in the *rlpi* package, while option *b* is a novel exploration. Note that the confidence intervals for all options are multiplicative and increase in width over time as the uncertainty of previous years is inherited by the rest of the trend.

### (iii) Time series length, number of data points and completeness

C-LPI data were collected from a variety of sources, which report on diverse species and geographies and are collected via different methodologies. Consequently, population time series vary in the number of data points (total number of years with non-null values), length (duration of time series from first to last year with non-null values), and completeness (number of data points divided by time period covered). In addition, there is no extrapolation beyond the start and end point of a time series, and thus each population time series enters the C-LPI at different points in time. Previous research has suggested that lower-quality population time series (shorter length, few data points and limited spatial coverage) diverge from average trends in higher-quality data and often exhibit more negative indices (Toszogyova *et al*. 2024; Marconi *et al*. 2021). The C-LPI requires three data points between 1970 and the end point of the index (currently 2022) but does not set criteria for length or completeness.

Thus, we explore three alternatives for the required number of data points:

a. Requirement of two or more data points as the global LPI approach of the *rlpi* package (Freeman *et al*. 2017) in alignment with Collen *et al*. 2009.
b. Requirement of six or more data points, in recognition that generalized additive models (GAM) can be applied to population time series with six or more data points, compared with those with fewer points (where instead, log-linear interpolation or linear regression are adopted; Marconi *et al*. 2021; Collen *et al*. 2009; Loh *et al*. 2005) (see section (iv) Modelling of short time series below).
c. Requirement of 15 or more data points to showcase the impact of longer time series.

Additionally, as there are no set criteria for length or completeness, we explored:

a. Four time series lengths (duration of time series from first to last year with non-null values): i) ≥5 years, ii) ≥10 years, iii) ≥15 years, iv) ≥20 years.
b. Four time series completeness levels: i) >0%, ii) >25%, iii) >50%, and iv) >75%.

### (iv) Modelling of short time series

Changes in population abundance are calculated by the geometric mean of relative abundance from 1970 to 2022, though modelling approaches to interpolate annual values between the start and end point of a time series differ, depending on the number of data points. In alignment with the global LPI (Collen *et al*. 2009), we used Generalized Additive Models (GAMs) to allow for a smoothed trend, rather than linear format (Wood 2017; Buckland *et al*. 2005) for population time series with six or more data points. GAMs permit interpolation for all years between the start and end years of the population time series. However, GAMs are less efficient with fewer data, and in some cases are not appropriate. Consequently, we apply alternative approaches for population time series with less than six data points or those with poor GAM fitness. The C-LPI uses linear regression models for time series with less than 6 data points. Here, we explore two alternatives for population time series where GAMs are not applied:

a. Log-linear interpolation, which is the global LPI default approach of the *rlpi* package (Freeman *et al*. 2017) in alignment with Collen *et al*. 2009 and Loh *et al*. 2005.
b. Forcing a GAM on short time series, as a numeric requirement for data points has not been identified.

### (v) Removal of outliers

While a geometric mean of relative abundance is considered a suitable and sensitive metric to assess biodiversity change (Santini *et al*. 2017; van Strien *et al*. 2012), it is nevertheless sensitive to outliers, which can be problematic if not addressed (Ledger *et al*. 2023; Leung *et al*. 2020; Buckland *et al*. 2011). The C-LPI does not currently remove outliers beyond the capping that’s currently built into the *rlpi* package. This step limits interannual change values (i.e., lambdas) to 1/-1, correspontherefore representative of the data includedding to a 10-fold increase or decline in abundance since the previous year — which recognizes large changes, but constrains their extent to a biologically plausible threshold. Here we explore three alternatives with removals of the most extreme logged interannual change values (species lambdas) on either side of the distribution (lower and upper extremes) at equal intervals:

a. Removing 5%.
b. Removing 10%.
c. Removing 15%.

We also explore removal of species lambda outliers from only the lower or upper extremes of the index for investigative purposes in alignment with Leung *et al*. (2020; see Supplementary Material) but caution that the removal of only extreme positives or declines introduces substantial bias for the interpretation of results.

### (vi) Weighting species

The data underlying the C-LPI are the result of 10 years of iterative data collection and cover approximately half of native vertebrate species (CESCC 2022) in Canada. The dataset is still continuously augmented, improving the taxonomic and spatial representation of the index for each subsequent iteration. However, gaps for species, habitats and geographies remain and should be accounted for to address potential biases of the collected data. Tackling taxonomic and ecosystem bias in the data through diversity weighting is an attempt to improve the representativeness of trends in the indicator (McRae *et al*. 2017) but can also reinforce potential biases within the dataset (e.g., where species with poor-quality data are weighted more heavily within the index) — essentially solving one problem, but potentially introducing another. The C-LPI weights all species equally within the index and is therefore representative of the data included.

a. Here we explore one alternative to an unweighted index that includes proportional weighting by species richness. The proportion of known native species was taken from the 2020 Wild Species Report (CESCC 2022) and used to assign a weight for each taxon (Table S1). This approach differs to McRae *et al*. (2017), as we did not include a secondary weighting of equal distribution among systems (i.e., freshwater, marine and terrestrial) for two reasons: (i) there were limited data for some taxonomic/systems groups such as marine herptiles, where data was only available for one species, and (ii) a lack of ecological rationale for equal weightings among systems.

### (vii) Baseline year

The baseline year influences the final calculation of the geometric mean. Importantly, the C-LPI (and the global LPI) is a relative measure, comparing trends to the baseline year of 1970. Therefore, any shift in the baseline year may result in differing overall average trends. Critically, 1970 has the fewest time series records, and thus selection of a year with more data records could theoretically be more accurate. Yet, shifting the baseline has an impact on the temporal length of the index and the longer-term conclusions that can be drawn. Final index values are reported relative to the baseline year.

a. Here, we explore differences in indices and average trends associated with shifting the baseline year at five-year increments from 1970 to 2015, while retaining the current dataset (i.e., time series that span the chosen baseline will still contribute to the index with the interpolated and actual data points available from the chosen baseline onwards).

## Interpretation and comparison

All indices use a benchmark value of 1 in 1970 (except for baseline year analysis), which is represented by a solid black line in the following graphs. A final index value >1.05 would represent an increase in average population abundance relative to 1970, while a value <0.95 would be representative of a decrease. Any value within ±0.05 of the baseline is considered stable (within 5%; Deinet *et al*. 2024). If confidence intervals are fully above or below the benchmark value of 1, we are more confident in the trajectory of the trend.

To compare outputs across methodological variants, we adopt a descriptive framework to evaluate sensitivity of the index, considering robustness as qualitative concordance in endpoints and trajectories across different methodological decisions. Specifically, we (i) report each index’s end-point value in 2022 relative to the baseline; (ii) summarize noteworthy differences in end-point values between variants; and (iii) assess trajectory consistency by visually inspecting overlaid time series, noting concordant directionality and any periods of divergence. Because there is no agreed quantitative benchmark for the ‘true’ trajectory of biodiversity trends, and our goal is transparency and accountability in indicator reporting rather than hypothesis testing, we do not perform formal statistical tests of between-variant differences. We emphasize end points because they reflect current value used in reporting, while qualitatively noting trajectory differences where they are noteworthy.

## Results

### (i) Treatment of zero values

The current C-LPI approach to treating zeros (replacement with NA) resulted in a trend and final index value (2022 index value = 0.90) in the midrange of values for the six alternative options explored (2022 index range = 0.81–0.98; Figure 1). Adding a small value of 0.000001 to zeros generated the greatest uncertainty around the LPI trend (2022 confidence interval range = 0.37), a contrasting trajectory (i.e., temporal trend) and the most positive final index value (2022 index value = 0.98). Alternatively, conditional replacement of leading and middle zeros with NA and treating trailing zeros by adding 1% of the mean of the population time series to all values within the time series containing zeros, resulted in the most negative trajectory and final index value (2022 index value = 0.81). Zero values were most common within time series (e.g. middle zeros) for fish, herptiles and marine mammals; however, trailing zeros were more common for terrestrial mammals (Figure S1). Within the C-LPI dataset, the proportion of zero values per population time series ranged from 0 to 0.85 (mean=0.02). Of these zero values, 65.2% were mammals, 32.2% fish, and 2.6% herptiles, while birds contained none as birds are represented in the dataset by long-term modelled indices from Environment and Climate Change Canada (ECCC).

**Figure 1.**
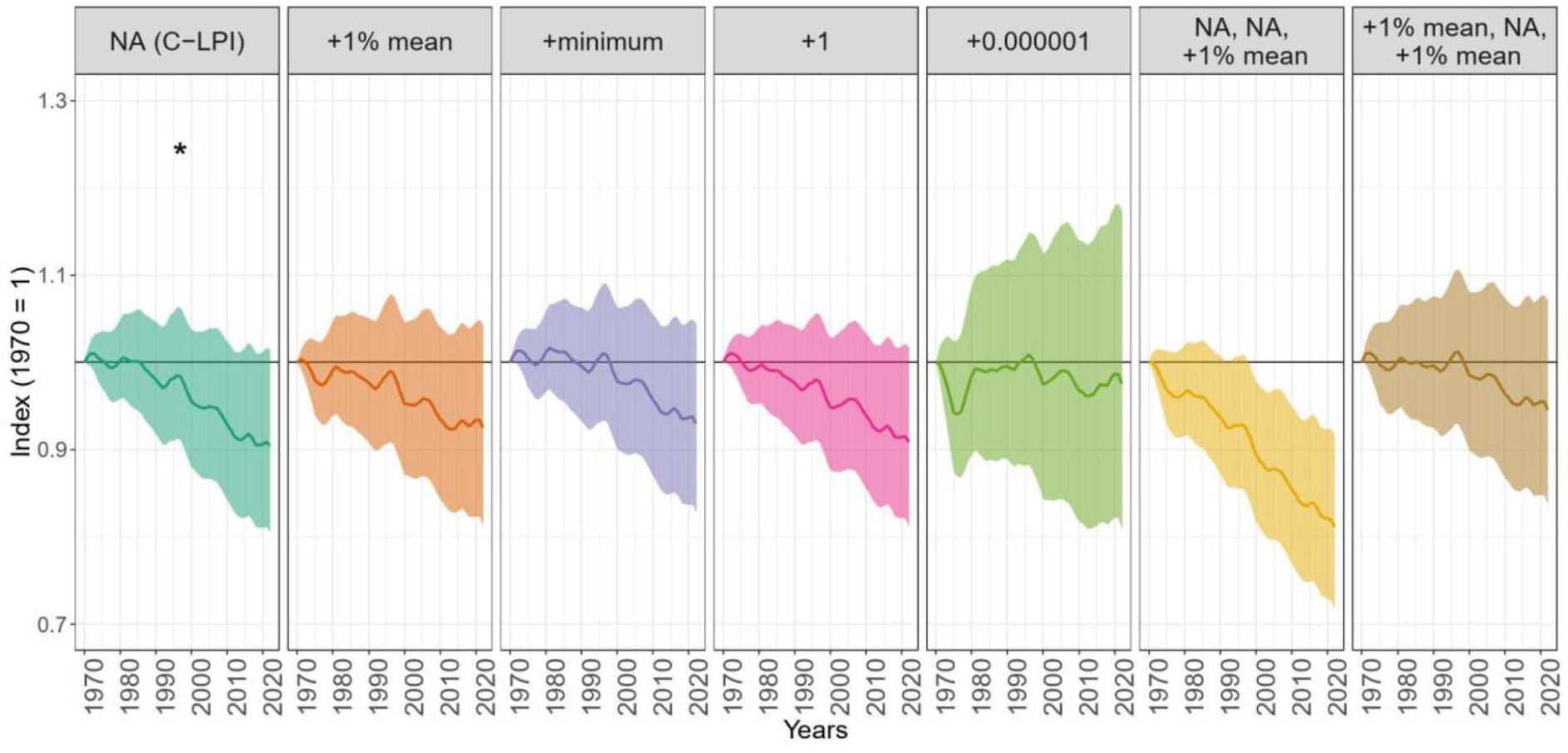
Evaluating the influence of different treatment options for zeros on the C-LPI trend, as indicated by banner labels. NA (C-LPI): zeros are treated as missing values (NA), current approach (indicated by an asterisk). Alternative methodological approaches include: (a) adding 1% of the mean of the population time series to all values within the time series containing zeros, (b) adding the minimum value of a time series to all values within the time series containing zeros, (c) adding 1 to all values within the time series containing zeros, (d) adding a very small number (0.0.000001) to zeros. Zeros may also be treated conditionally depending on their position in the time series:(e) leading, middle, and trailing zeros may be replaced respectively with NA, NA, and 1% of the mean of the population time series to all values within the time series containing zeros, or (f) with 1% of the mean of the population time series to all values within the time series containing zeros, NA, and 1% of the mean of the population time series to all values within the time series containing zeros.

### (ii) Confidence intervals and uncertainty

Bootstrapping by year resulted in the smallest final confidence intervals (2022 confidence interval range = 0.85–0.96) surrounding the C-LPI, corresponding to minimal variability among confidence interval values relative to bootstrapping by population or species (Figure 2). Alternatively, the current C-LPI approach (bootstrapping by species) showcased the largest final confidence intervals (2022 confidence interval range = 0.80–1.02, marginally larger than bootstrapping by populations), providing greater transparency in the range of potential index values.

**Figure 2.**
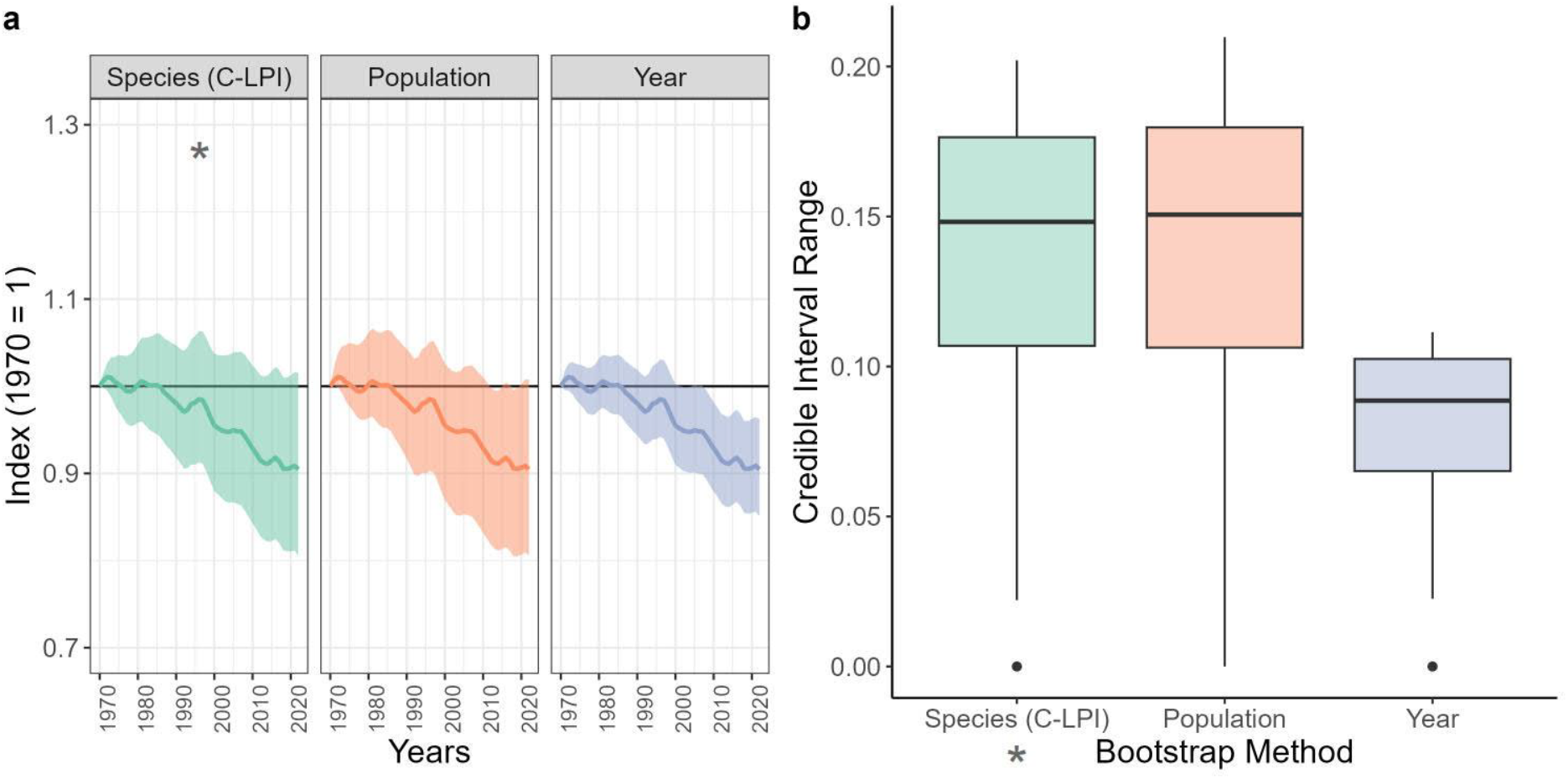
Differences in confidence intervals bootstrapped via three different methods (population, species, and year bootstrap), showcased via the (a) index, and (b) confidence interval range calculated per year. Note that the current C-LPI approach adopts species bootstrapping (indicated by an asterisk).

### (iii) Time series length, number of data points and completeness

The current C-LPI approach, which includes only time series with ≥3 data points, results in a declining temporal trend with a final index value of 0.90, and is within the midrange of the alternative three options explored (2022 index range = 0.85–1.03; Figure 3). Permitting ≥2 data points generated the greatest declining trajectory and final index value in 2022, ≥6 data points generated an upward temporal trend and final index value while restricting to ≥15 data points resulted in the most similar final index value to the C-LPI (though it showcases a slightly different trajectory, initially increasing and only falling below the baseline beginning in 2010). This is intriguing given that more than half of the population time series data is eliminated to achieve the requirement of ≥15 data points (Table S2).

**Figure 3.**
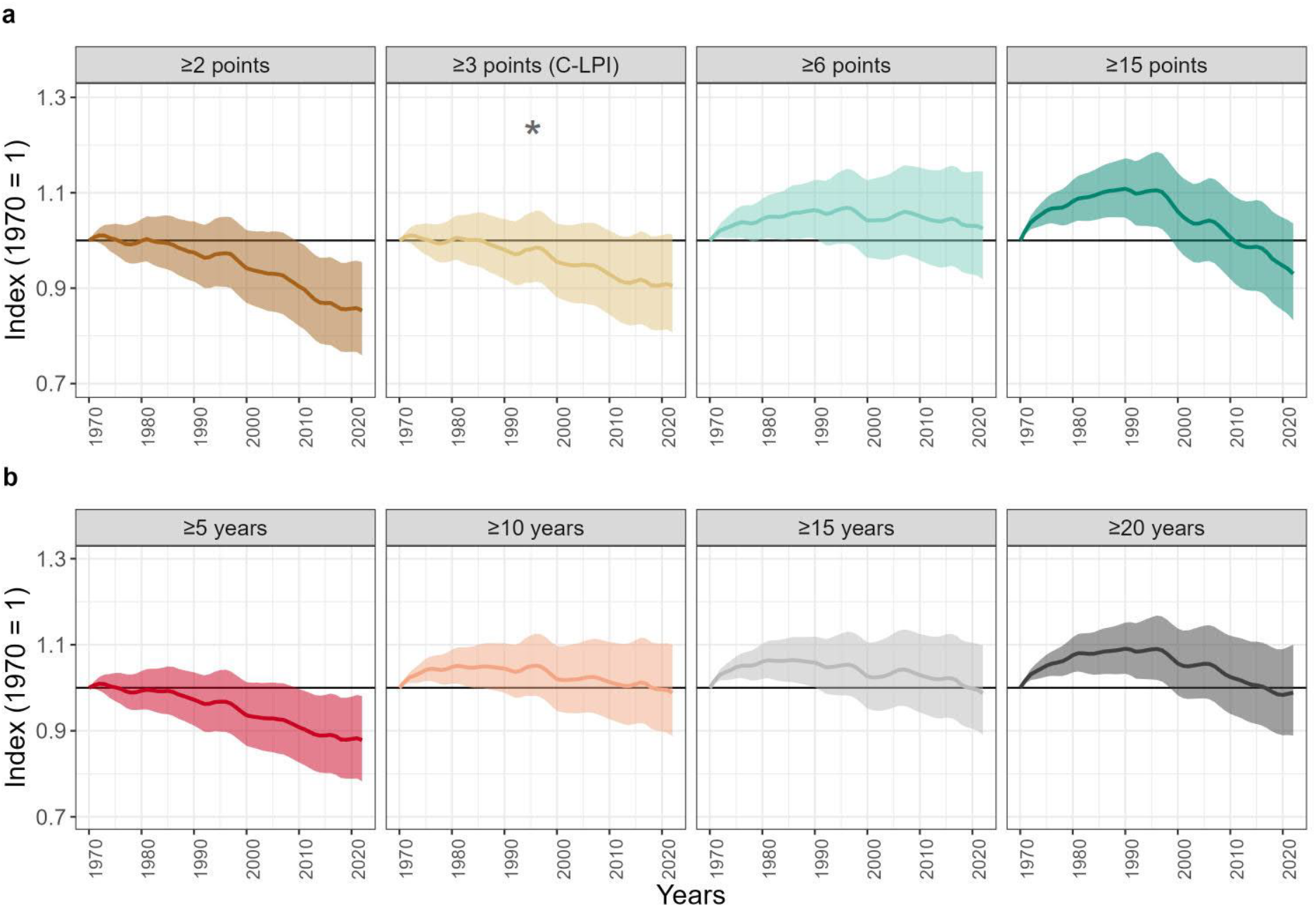
Evaluating the differences between data inclusion criteria on the index, pertaining to (a) number of data points and (b) length (time period covered) for population time series. Note that the current approach adopted by the C-LPI (indicated by an asterisk) requires a population time series with ≥3 data points but has no set criteria for time series length. As the criteria for data inclusion differs among these options, so do the number of species and populations included within each index.

Alternatively, time series length shows a range of final index values from 0.87 to 0.99 for the four options explored, where inclusion of shorter time series length produces a more negative trajectory and final index value relative to indices that contain data restricted to longer time length (Figure 3). Given the substantial overlap in confidence intervals, there were, however, no noteworthy qualitative differences among trends with ≥10, ≥15, and ≥20 years, despite changes in the number of species included (825, 733 and 631, respectively). An assessment of completeness (number of data points divided by duration of time series from first to last year with non-null values), reveals greater variation among indices that include incomplete time series relative to those restricted to a ≥50% completeness level (Figure S2). Final index values range from 0.82–1.02 for the 16 completeness permutations, with considerable variability among temporal trends.

### (iv) Modelling of short time series

Approximately 36% (n=1842) of population time series were modelled using a GAM, meaning that 64% (n=3257) of time series (i.e., <6 data points or poor GAM fitness) required alternative methods for modelling. Adoption of linear regression for these latter time series (2022 index value =0.90) resulted in only minor differences relative to forcing all trends through a GAM (2022 index value = 0.89) (Figure 4). Unsurprisingly, log-linear interpolation resulted in more interannual variability, and also exhibited a slightly more positive final index value of 0.95. Nevertheless, all modelling approaches resulted in a declining trajectory relative to 1970.

**Figure 4.**
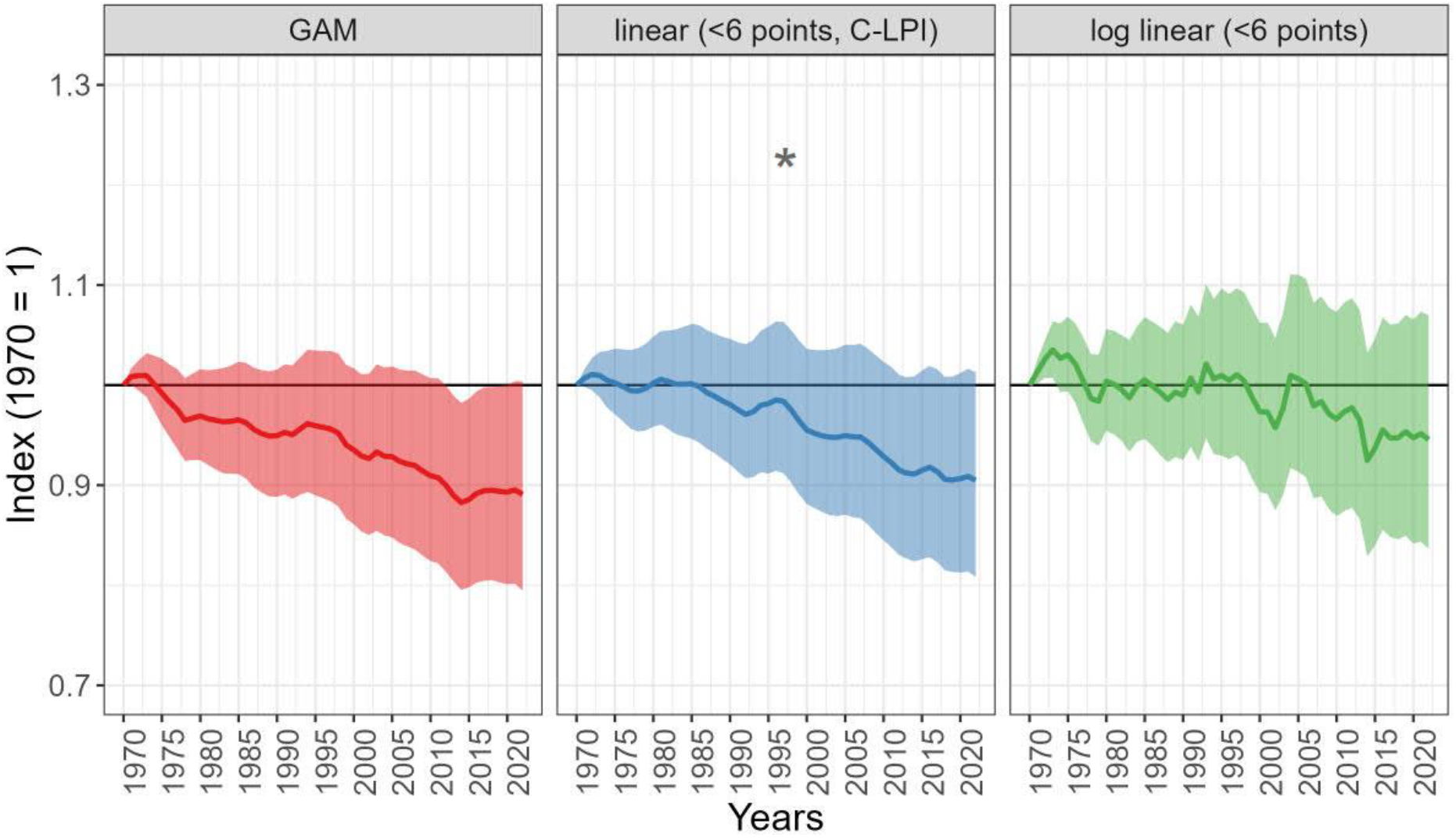
Comparison of indices using differing modelling approaches, where Generalized Additive Models (GAM) are used for population time series with ≥6 data points (n = 1842), combined with alternative modelling approaches for short time series or those that resulted in a poor GAM fit, including (a) forcing a GAM (n = 3257), (b) applying linear regression models (n = 3257), and (c) use of log-linear interpolation (n = 3257). Note that the linear regression method is currently adopted by the C-LPI, as indicated by an asterisk.

### (V) Removal of outliers

The removal of species lambda outliers from both upper and lower extremes of the index resulted in progressively smaller confidence intervals but similar values, with a noteworthy decline in the late 1990s (i.e., differing temporal trajectory; Figure 5). The C-LPI does not currently remove outliers and exhibits a final index value of 0.90. Removal of 15% of outliers from both the upper and lower extremes results in a final index value of 0.91, albeit with smaller confidence intervals, approximately 49% narrower compared to not removing outliers. Separate removal of upper and lower extreme outliers has substantially larger impacts on LPI trends and final index values, given the bias introduced to the dataset. There is greater variability in the trajectories and final index values of indices when the most declining trends are removed, relative to the removal of the most increasing trends (Figure S3).

**Figure 5.**
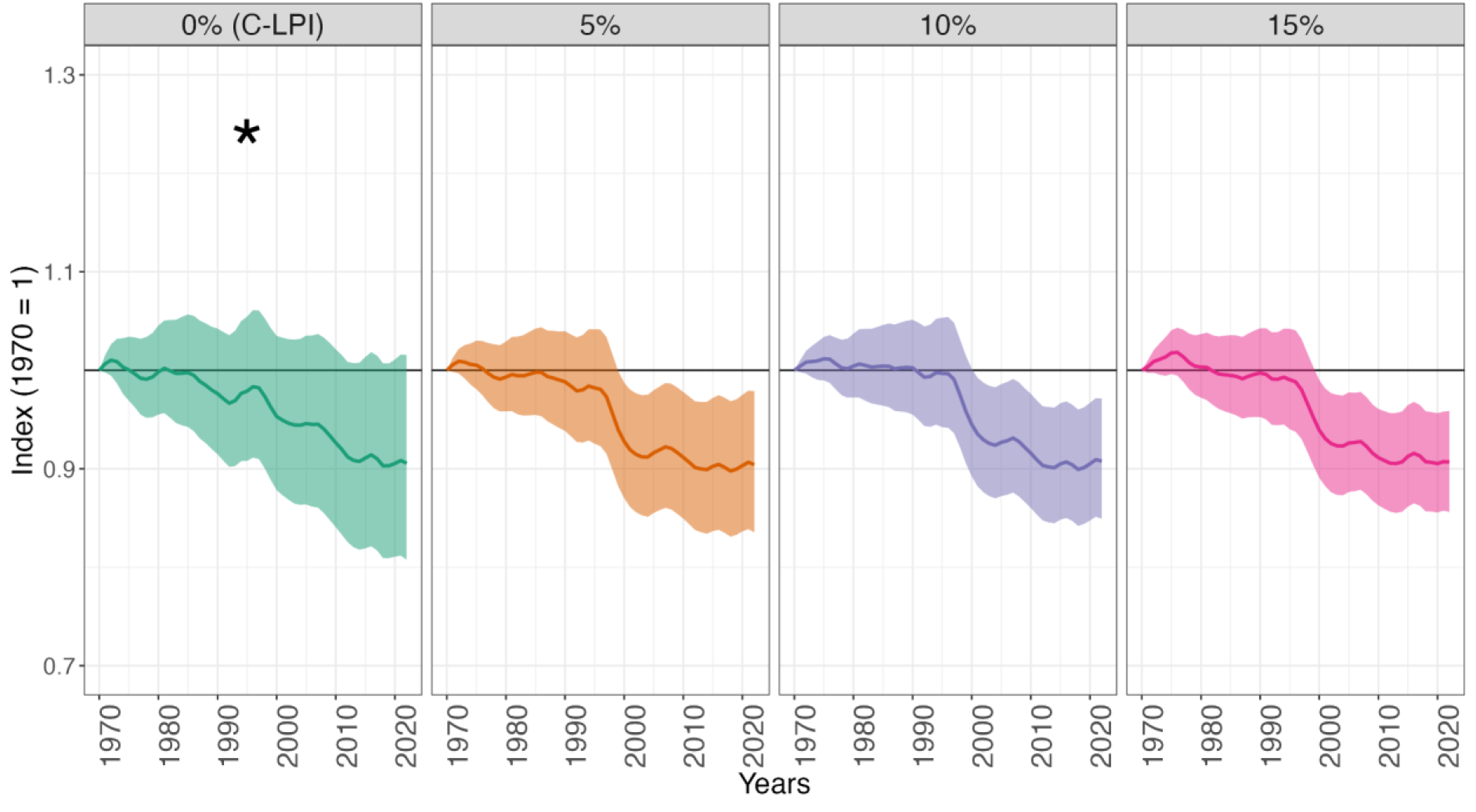
Removal of 0%, 5%, 10%, and 15% of species lambda outliers from both the lower and upper extremes of the index. Note that outliers are currently not removed in the C-LPI (i.e., 0%), as indicated by an asterisk.

### (vi) Weighting species

The weighted index shows greater temporal variability from 1970-2022, but both approaches result in similar final index values (2022 weighted index = 0.91 vs 2022 unweighted index = 0.90; Figure 6).

**Figure 6.**
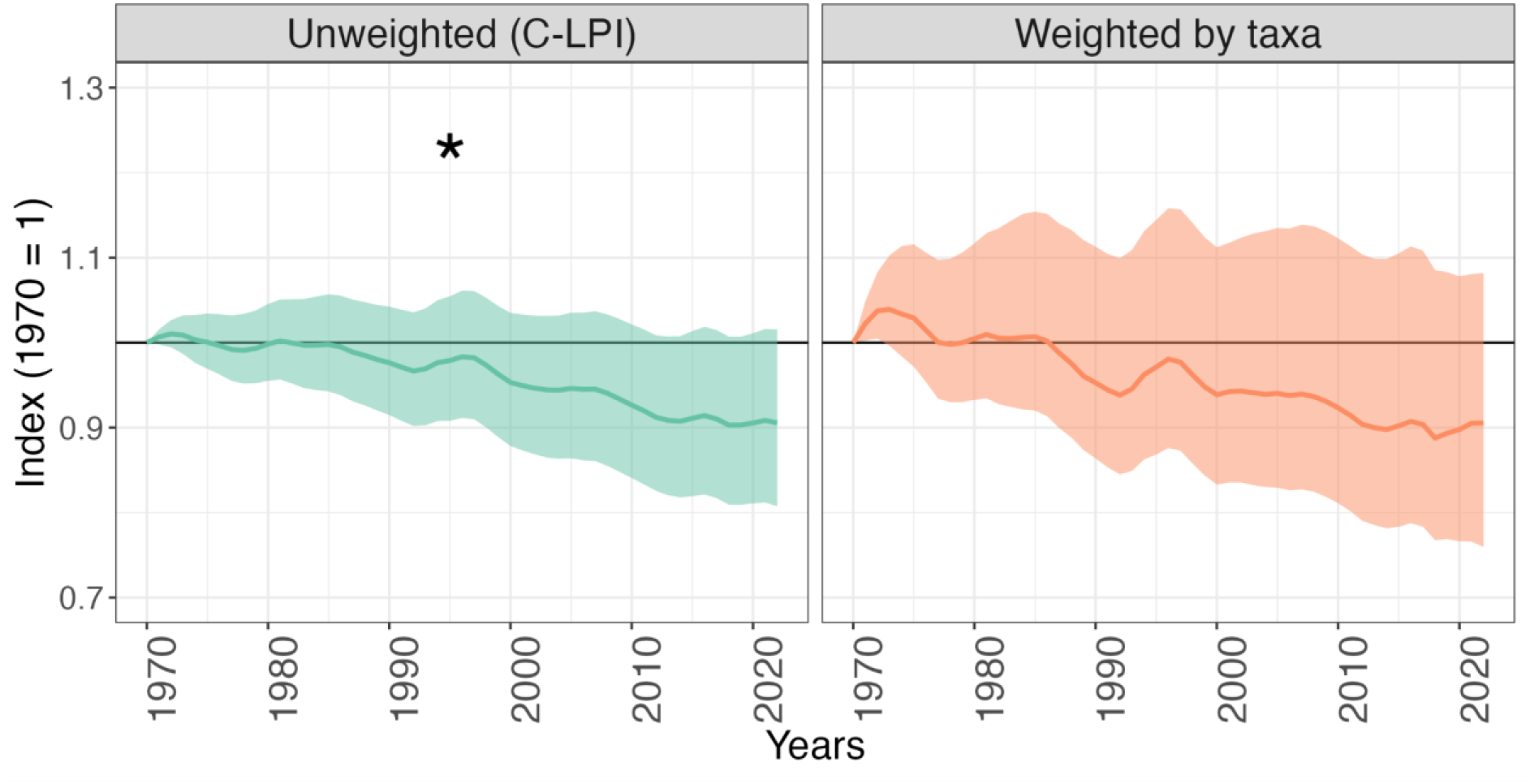
Lambda values per taxonomic group (i.e. birds, mammals, herptiles, and fish) (a) contributed equally to the final index value in the unweighted trend (current C-LPI approach indicated by an asterisk) or otherwise were (b) weighted proportional to species richness of each taxonomic group in the weighted approach.

Notably, the weighted index yields larger confidence intervals (2022 confidence interval range = 0.76– 1.08) relative to the unweighted approach (2022 confidence interval range = 0.80–1.02).

### (vi) Baseline year

Indices exhibited relatively similar trends despite shifting the baseline year, with final index values ranging from a low of 0.90 using a baseline of 1975 to a high of 0.99 using a baseline of 2015 (Figure 7). The distribution of yearly index values by baseline year exhibited a skewed temporal pattern, with highest values occurring when the baseline year was closest to present day (Figure 7). There was less variation among indices and average lambdas when selecting more recent baseline years—due to limited temporal extent (Figure 7). The cut-off year for different baselines was applied to the modelled data so that time series that spanned the chosen baseline still contributed to the index with the interpolated and actual data points available from the chosen baseline onwards. The dataset for the index with a baseline in 1970 yielded the greatest number of population time series, whereas the baseline of 2015 contained the least (Table S3).

**Figure 7.**
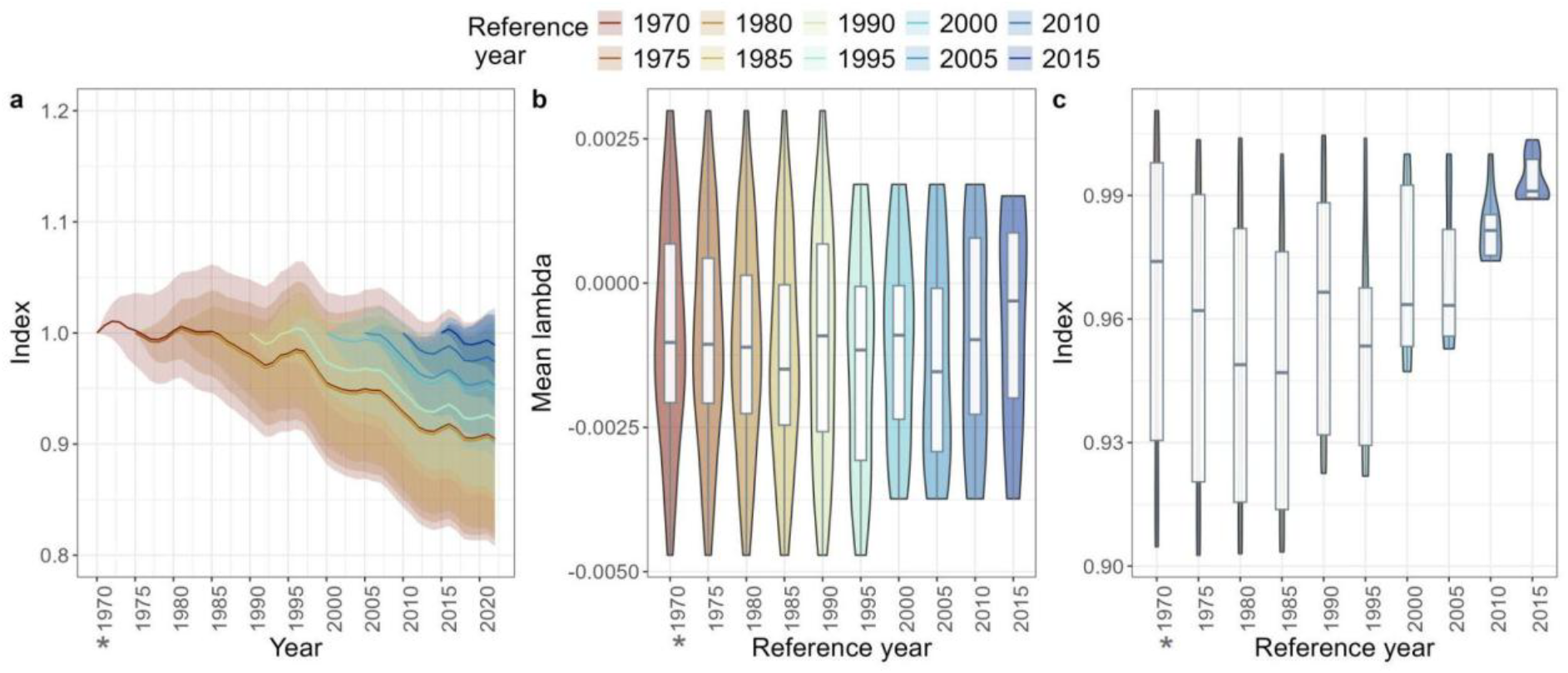
Comparison of indices using baseline years that shift by 5-year increments, as depicted through (a) indices, (b) average yearly lambda values, and (c) yearly index values. Note that the C-LPI uses 1970 as the baseline year (as indicated by an asterisk). The number of populations and species contributing to each baseline calculation are shown in Table S3.

## Discussion

We evaluated the influence of various decision points in the analysis pipeline of the Canadian Living Planet Index (C-LPI). Within our analysis, index endpoints were relatively similar for most decision points. Assessments pertaining to the treatment of zeros, number of data points and completeness contributed the greatest variability in trends, spanning final index values of 0.81–0.98, 0.85–1.03, 0.82-1.02, respectively (i.e., up to a 20% difference). Notably, however, these assessments also adopted some of the most permutations for investigation (7, 4, and 16, respectively). These remain challenges for the LPI, given that population time series data varies considerably in length and number of data points (Dover *et al*. 2023; Ledger *et al*. 2023), and zero values are not tolerated in the calculation of the geometric mean. By contrast, there was greater variability within index trajectories, which may have implications for the interpretation of the index and potential conservation actions. Notably, the C-LPI approach frequently generated a trend and final index value that was in the mid-range of methodological options and permutations explored (i.e., rarely an outlier), which is a promising future benchmark for evaluation.

### (i) Treatment of zeros

The most common approach to address zeros is to add a small number to zeros to permit mathematical calculations (Santini *et al*. 2017; van Strien *et al*. 2012; Buckland *et al*. 2011; Collen *et al*. 2009; Buckland *et al*. 2005). However, we provide — for the first time — an in-depth analysis of zero location within time series, and upon manual inspection of the data, find that zero values rarely reflect local extirpations in the C-LPI. Instead, zeros are commonly representative of missing values or failure to detect a species within the confines of the study year. In the C-LPI, zeros were most common within population time series for fish, herptiles and marine mammals (Figure S1). Trailing zeros — which may be more reflective of local extirpation — were more common among terrestrial mammals. However, trailing zeros for terrestrial mammals were only included in 20 out of hundreds of data sources, and were most common for widespread species, including black bear (*Ursus americanus*; Garsehlis *et al*. 2016) and red fox (*Vulpes vulpe*s; Hoffman and Sillero-Zubiri 2021), or those with well-documented boom and bust cycles (*Lepus americanus*; Krebs *et al*. 2001), suggesting that these zeros were more reflective of missing observations or low detectability, rather than extirpation. In line with the underlying data and recent recommendations (Toszogyova *et al*. 2024), the C-LPI treats zeros as missing values. To our knowledge, our study was the first instance exploring conditional treatment of zeros within population time series. Interestingly, conditional treatment of zeros resulted in the greatest divergence from both the baseline and alternative approaches. However, this approach may only be suitable with an in-depth knowledge of where and why zero values are recorded and may not be amenable to broader-scale analyses.

### (ii) Confidence intervals and uncertainty

The confidence interval approach adopted by the C-LPI comprises the greatest range of possible values. This approach not only showcases a broader possible range of values for improving transparency and interpretation of the index but can also help inform suitable conservation actions by assessing whether there are broad differences among species (Rowland *et al*. 2021), and whether species-specific actions may be more appropriate.

### (iii) Time series length, number of data points and completeness and (iv) Modelling of short time series

Some species and regions are less well-studied, resulting in shorter and sparser time series (Marconi *et al*. 2021). While time series with fewer data points may be informative and representative of true patterns, they can introduce bias into the index because of the underlying mathematical calculations, particularly if they are not reflective of the broader geographical range of a species. Not only are short time series on average declining (Toszogyova *et al*. 2024; Marconi *et al*. 2021), but they are treated differently within the modelling of the index, as they are processed by methods that do not adopt data smoothing (note that longer time series — those with six or more data points — employ a GAM method). Consequently, stochasticity and uncertainty are retained, which can further impact biases of short and sparse time series (Toszogyova *et al*. 2024; Hébert & Gravel 2023). Notably, however, options for modelling of short time series in our analysis, yielded only a 6% range in the final index values produced, suggesting that individually modelling of short time series may not bias the C-LPI, but when combined with other methodological decisions, the impact could be greater. In the absence of longer, fuller and more complete data on which a GAM can be applied, transparency on data gaps and coverage is essential for more appropriate interpretation of results.

### (v) Removal of outliers, (vi) Weighting species and (vii) Baseline year

Despite the discourse in the literature surrounding extreme increases and decreases and their effect on the overall indicator (Toszogyova *et al*. 2024; Leung *et al*. 2022b; Murali *et al*. 2022; Leung *et al*. 2020), we found that the removal of species lambda outliers from both the lower and upper extremes resulted in the least variability in final index values (i.e., end points) among the various methodological analyses explored (range: 0.90–0.91), though differences among trajectories are more substantial. Discrepancies from global analyses may be due, in part, to the removal of zero values (i.e., treating zeros as NAs), as well as an equal weighting system, which is representative of the data included and limits the exacerbation of extreme trends. On the other hand, population abundance data for the Nearctic realm often exhibits relatively stable trends, and thus the impact of outliers may also be less drastic (Leung *et al*. 2020). Similarly, shifting baselines seemed to have relatively minimal impact on final index values, likely because the interpolated data points were retained in our analysis to inform a longer-term trend. Nevertheless, it’s important to remember that the LPI is multiplicative, and thus extreme fluctuations at the beginning of the study period are transferred down, irrespective of when the baseline is set (Toszogyova *et al*. 2024). Outliers can also have compounding effects with other methodological choices, such as species-richness weightings, where extreme increases or declines are further emphasized through weighting a species more heavily. However, contrary to the global LPI (McRae *et al*. 2017) — although calculated differently — species-richness weightings did not substantially impact final index values, in part, because the Canadian dataset is more robust — representing half of native vertebrate species.

## Recommendations

Within the C-LPI, there are no perfect methodological choices — each has contrasting evidence and opinions, with corresponding benefits and drawbacks. By showcasing trends and options more transparently, we provide a greater array of information to accompany the temporal trend (i.e., trajectory) and final index value (i.e., end point) of the C-LPI. Importantly, the intent of our research was not to identify the “best” approach, but rather to provide transparency and accountability in reporting. Nevertheless, our research identifies three methodological decisions that produce variation in final index values up to 20% (number of data points, treatment of zeros and data completeness) which should be the subject of further research development and scientific consensus to reduce the variability within index end points and trajectories.

Our research contributes to a growing portfolio that reinforces the robustness of monitored vertebrate abundance data in Canada. However, we call for continued investment in long-term biodiversity monitoring to fill outstanding data gaps and address biases in data collection (Toszogyova *et al*. 2024; Leung *et al*. 2022b; Murali *et al*. 2022) and to continue investigating biodiversity trends to improve efficacy of the C-LPI and associated analyses that rely on similar data. Broader geographic coverage will be important for capturing localized dynamics and differences among populations of the same species (Marconi *et al*. 2021). For instance, wolverines have declined in southern and eastern Canada but have exhibited more stable trends elsewhere (COSEWIC 2014). Geographic coverage extending to taiga and Arctic ecozones may also be particularly important due to the disproportionate effects of climate change in higher latitudes (IPCC 2021). Moreover, previous studies have suggested that data monitoring and collection in Canada be targeted towards small fishes and mammals, as well as carnivorous reptiles to enhance the representation of biotic traits (Currie *et al*. 2022). Continued and expanded monitoring will be key to appropriately evaluating trends in monitored vertebrate abundance. In addition, analyses at taxonomic or system-level can support greater investigation and further unpack the status of potential vulnerable groups.

Even though there are criticisms of the LPI as a global indicator (Johnson *et al*. 2024; Toszogyova *et al*. 2024; Leung *et al*. 2022a; Leung *et al*. 2022b; Leung *et al*. 2022c; Loreau *et al*. 2022; Murali *et al*. 2022; Puurtinen *et al*. 2022; Buschke *et al*. 2021; Leung *et al*. 2020), they do not diminish the fact that global biodiversity is in peril (Toszogyova *et al*. 2024; IPBES 2018). The criticisms of the LPI should not detract from the urgency of addressing biodiversity loss, but they highlight the complexities of measuring, interpreting and understanding trends in biodiversity. As Parties to the Convention on Biological Diversity (CBD) continue to develop, report on and implement National Biodiversity Strategies and Actions Plans (NBSAPs), component (i.e., optional) indicators such as the LPI are important for broadening the coverage of the Monitoring Framework of the GBF, and can help to address gaps (Affinito *et al*. 2024). Canada boasts the best representation for a national indicator (Marconi *et al*. 2021), which not only strengthens the utility of the indicator at a national level but allows for greater exploration of the data and methodological choices while avoiding some of the criticized issues. We recognize that Canada is in a privileged position (Zhang *et al*. 2023), and that only a subset of Parties to the CBD may be able to replicate our methodological analysis with a similar robustness of associated outputs—nevertheless, we encourage the uptake of component indicators in NBSAPs, and recommend broader transparency of methodological decisions for all applications and disaggregations of indicators, such as the LPI. It’s imperative that biodiversity indicators are clearly communicated, transparent and simple to understand in order to appropriately operationalize their use in decision-making (McQuatters-Gollop *et al*. 2019) and support policies and conservation actions that are strategically sound.

This study adds to a growing portfolio that reinforces the strength of monitored vertebrate abundance data in Canada (Currie *et al*. 2022; Marconi *et al*. 2021), supporting its utility as a national biodiversity indicator. We hope that the transparency provided in this study will further strengthen the utility of the C-LPI and its outputs, given its use as an adopted biodiversity indicator for domestic reporting on the GBF. It is unrealistic to assume that we can feasibly measure population trajectories of all species in all geographies across large temporal extents, and this reality necessitates decision-making on imperfect data. It is therefore vital to provide transparency in the methodological decisions that underpin biodiversity indicators in order to provide decision makers with the necessary information to appropriately interpret patterns, evaluate progress, and inform conservation action.

## Acknowledgements

This manuscript is a product of a working group supported by WWF-Canada, the Zoological Society of London and the Living Data Project (LDP), an initiative of the Canadian Institute of Ecology and Evolution. The LDP is funded by a Collaborative Research and Training Experience (CREATE) grant from the Natural Science and Engineering Research Council of Canada (NSERC). The data from the Living Planet Index Database used in our analysis have been collected by countless individuals across the country. We thank them for their contributions over the years and for sharing their hard-earned data to make this work possible.

## Footnotes

1 The Canadian Species Index (CSI) is a domestic indicator in Canada’s National Biodiversity Strategy and Action Plan (NBSAP), known as the 2030 Nature Strategy. Previous iterations of the C-LPI (WWF-Canada 2020) and CSI (Environment and Climate Change Canada; ECCC 2023) had only minor methodological differences related to the criteria of data for inclusion (i.e., a requirement of three data points in the former and two data points in the latter). However, as of 2025, approaches have been harmonized across organizations, and thus, future discrepancies will be a result of new population time series data only.

2 Henceforth referred to as the C-LPI.

3 These data will be uploaded to the Living Planet Index Database in 2026

