## Supplementary Material for "Navigating methodological decisions: Balancing rigor and data volume of the Canadian Living Planet Index"

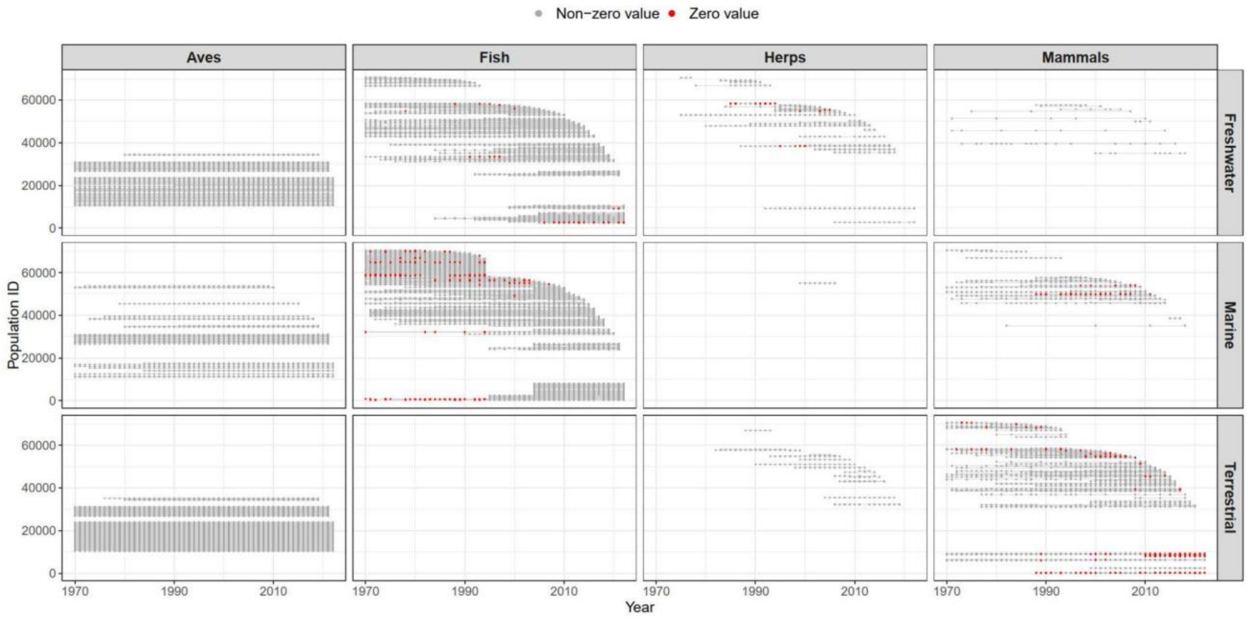


**Figure S1.** Location of zeros in each time series, subset by taxa and system, and ordered by time series end date. Only time series with ≥3 data points — those included in the C-LPI — are included.

**Table S1.** The relative weight of each taxon's contribution to the overall C-LPI trendline in the weighted trend analysis.

| Taxon | Weighting |
|---|---|
| Birds | 0.254218 |
| Mammals | 0.110236 |
| Herptiles | 0.048931 |
| Fish | 0.586614 |

**Table S2.** Number of species and population time series contributing to each trend for assessing the number of data points required and time series length.

| Variable | Criteria | Species | Population time series |
|---|---|---|---|
| Number of data points | ≥2 data points | 950 | 5793 |
| | ≥3 data points | 910 | 5099 |
| | ≥6 data points | 805 | 3612 |
| | ≥15 data points | 600 | 1502 |
| Length (time period covered) | ≥5 years | 901 | 4936 |
| | ≥10 years | 825 | 3835 |
| | ≥15 years | 733 | 2737 |
| | ≥20 years | 631 | 1642 |

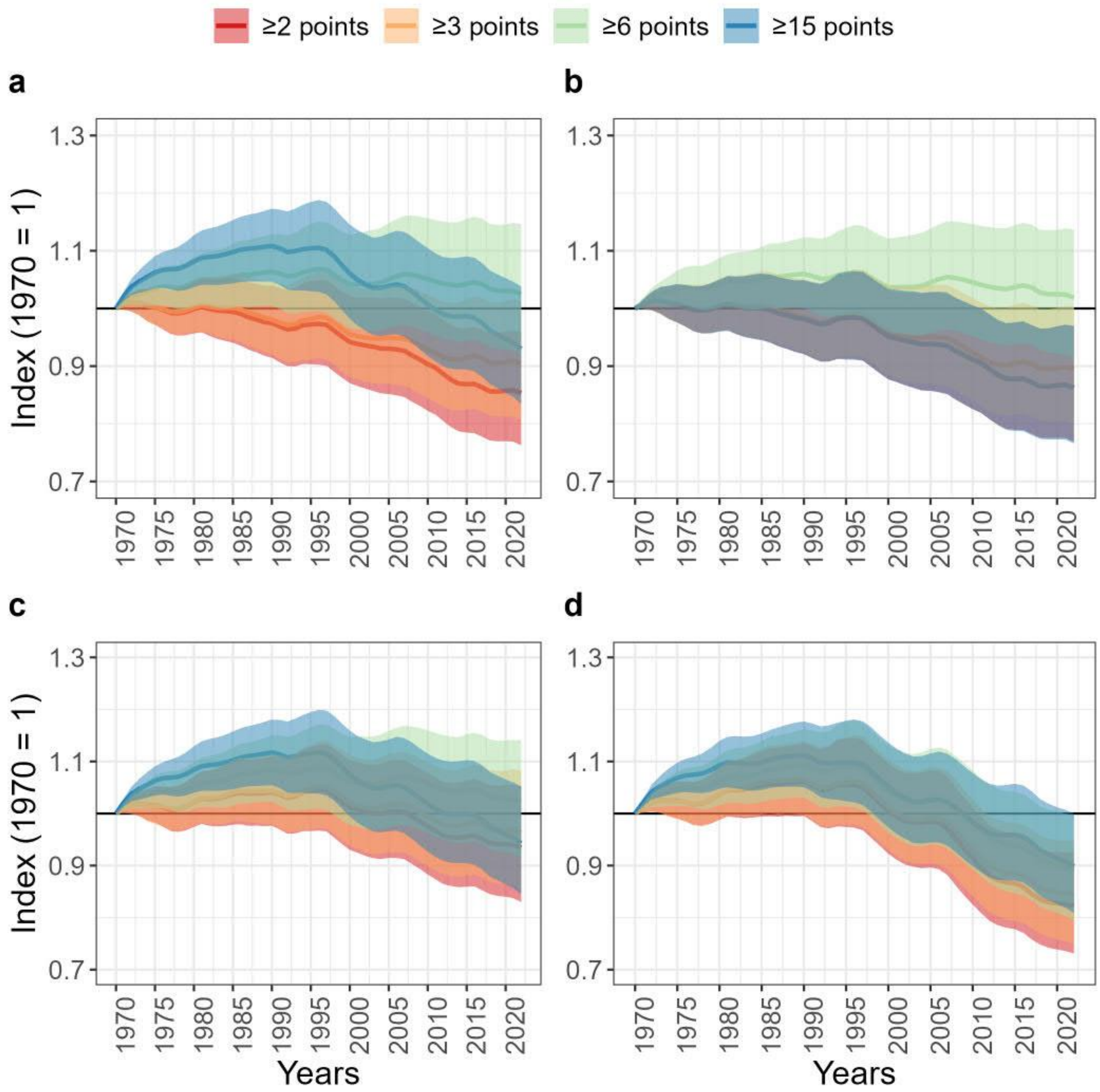


**Figure S2.** Evaluating the differences among the criteria for data inclusion on the index, as it pertains to population time series completeness at equal intervals of a (a) 0%, (b) 25%, (c) 50% and (d) 75% completeness score, for time series that contain ≥2, 3, 6, or 15 data points (recognizing that a 100% completeness score for 2 data points may differ significantly from 15 data points). Completeness is calculated based on the number of data points in a time series divided by the time period it covers (i.e., length), where, for instance, a time series from 2015-2020 with 2 data points is considered 40% complete). Note that there is no current requirement for time series length, nor completeness within the C-LPI. As the criteria for data inclusion differs among these options, so does the number of species and populations contributing to each index.

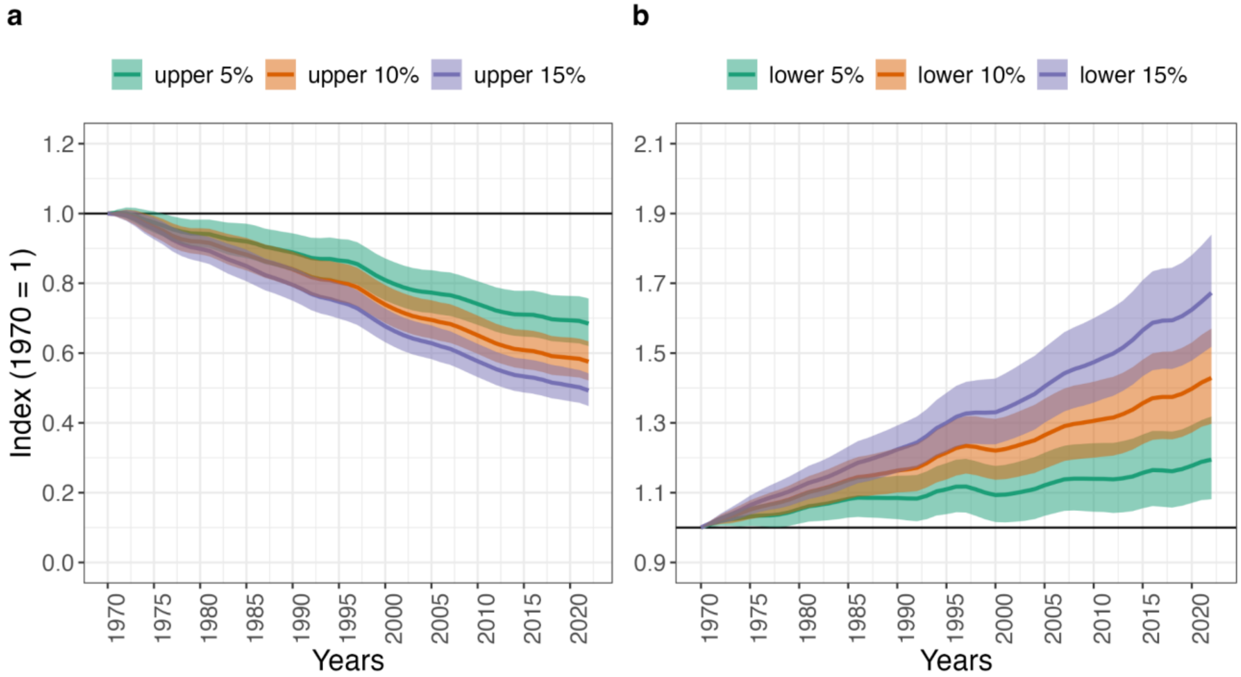


**Figure S3.** Independent removal of (a) upper and (b) lower lambda outliers, corresponding to 5%, 10%, and 15% extreme thresholds.

**Table S3.** Number of species and population time series contributing to each trend for assessing changes in indices with different baseline years. Note that time series with ≤2 non-NA data points are excluded from these counts, as per C-LPI inclusion criteria (Table 1).

| Reference year | Population time series | Species |
|---|---|---|
| 1970 | 5099 | 910 |
| 1975 | 5055 | 907 |
| 1980 | 4960 | 904 |
| 1985 | 4841 | 900 |
| 1990 | 4617 | 889 |
| 1995 | 4025 | 863 |
| 2000 | 3894 | 845 |
| 2005 | 3585 | 808 |
| 2010 | 2736 | 736 |
| 2015 | 1711 | 639 |
